# Resting-state fMRI data of awake dogs *(Canis familiaris)* via group-level independent component analysis reveal multiple, spatially distributed resting-state networks

**DOI:** 10.1101/409532

**Authors:** Dόra Szabό, Kálmán Czeibert, Ádám Kettinger, Márta Gácsi, Attila Andics, Ádám Miklόsi, Enikő Kubinyi

## Abstract

Resting-state networks are spatially distributed, functionally connected brain regions. Studying these networks gives us information about the large-scale functional organization of the brain and alternations in these networks are considered to play a role in a wide range of neurological conditions and aging. To describe resting-state networks in dogs, we measured 22 awake, unrestrained animals of either sex and carried out group-level spatial independent component analysis to explore whole-brain connectivity patterns. Using resting-state functional magnetic resonance imaging (rs-fMRI), in this exploratory study we found multiple resting-state networks in dogs, which resemble the pattern described in humans. We report the following dog resting-state networks: default mode network (DMN), visual network (VIS), sensorimotor network (SMN), combined auditory (AUD)-saliency (SAL) network and cerebellar network (CER). The DMN, similarly to Primates, but unlike previous studies in dogs, showed antero-posterior connectedness with involvement of hippocampal and lateral temporal regions. The results give us insight into the resting-state networks of awake animals from a taxon beyond rodents through a non-invasive method.

## Introduction

Resting-state networks (RSNs) are spatially distributed, functionally connected brain regions, characterized by the correlation of the time series of spontaneous, low frequency (0.01-0.1 Hz) fluctuations of the blood-oxygen level dependent (BOLD) signal, usually acquired in the absence of a specific task, and measured via functional magnetic resonance imaging (fMRI). The structure and assumed tasks of RSNs are of high interest as they have the potential to provide us novel insights and additional information about the brain’s large scale functional organization^1, 2^, and alternations in these networks have been found to correspond with various pathologies such as dementia or ADHD^3^.

As a result, the number of human rs-fMRI studies grew rapidly in recent years, but the evolutionary origin of these networks received less attention. To reveal phylogenetic changes and conserved core physiological mechanisms, it is crucial to compare a diverse range of non-human species^4^. To date, resting-state networks were investigated via fMRI in mice^5^, rats^6^, marmosets^7^ macaques^8^, dogs^9^ and ferrets^10^. Sensorimotor networks, such as visual and/or somatosensory networks have been described in most animal resting-state fMRI studies, but while in rodents there has been no report on a visual network separated from the somatosensory network, a visual network was described in ferrets^10^, while multiple visual networks were reported in primates^7^. Salience-like networks so far have been described in rodents^5^ and primates^7^, while fronto-parietal like components have been found only in primates^8^.

One widely used method to explore these whole-brain connectivity patterns is spatial independent component analysis (ICA), a data driven, model-free method^2^, appropriate to describe networks in case of a species which brain’s functional characteristics are yet to be determined, as it does not require selection of a priori seed regions. This method attempts to discover statistically independent source signals from the measured observations, using non-linear transformations while looking for spatial independence^11^.

Although putatuve default mode networks have been described in all of the investigated species, the typical antero-posterior connectedness of the human default mode network was only found in superorder Euarchontoglires (primates and rodents), while the corresponding networks in superorder Laurasiatheria (described in ferrets^10^ and dogs^9, 12^) were reported to show antero-posterior dissociation. We aimed to investigate whether and if so, what kind of spatially distributed resting-state networks are detectable in a larger sample of awake, unrestrained family dogs in a resting state fMRI setup, following up on previous reports^9,12^

## Results

Group-ICA decomposed the data into 20 independent components. ICASSO analysis returned a high stability index of the 20 estimate clusters (mean *±*SD = 0.940 *±*0.008), indicating high consistency across multiple ICA runs. Examination of the components resulted in retaining 6 components as signal components (spatially distributed networks with low frequency fluctuations, showing high correspondence with bilateral grey matter areas based on their spectral characteristics and spatial maps), while the remaining 14 components were classified as miscellaneous or noise. The excluded components, among others contained ventricular, susceptibility (located mainly in the frontal lobe, due to the extensive sinuses present in the dogs cranium) and motion artefacts. Signal components were primarily restricted to cortical areas. The resulting signal components were as follows:

RSN A (Fig. 1): This resting-state network covered parts of the gyrus compositus rostralis, rostro-dorsal regions of the gyrus cinguli and the gyrus rectus; medial, bilateral caudal regions of the gyrus cinguli and gyrus splenialis, included bilaterally the hippocampus and gyrus parahippocampalis, the gyrus compositus caudalis, and caudoventral regions of the cerebellum. All of these regions correspond to regions indicated in the default mode network (DMN) described previously in humans^13^.

**Figure 1.**
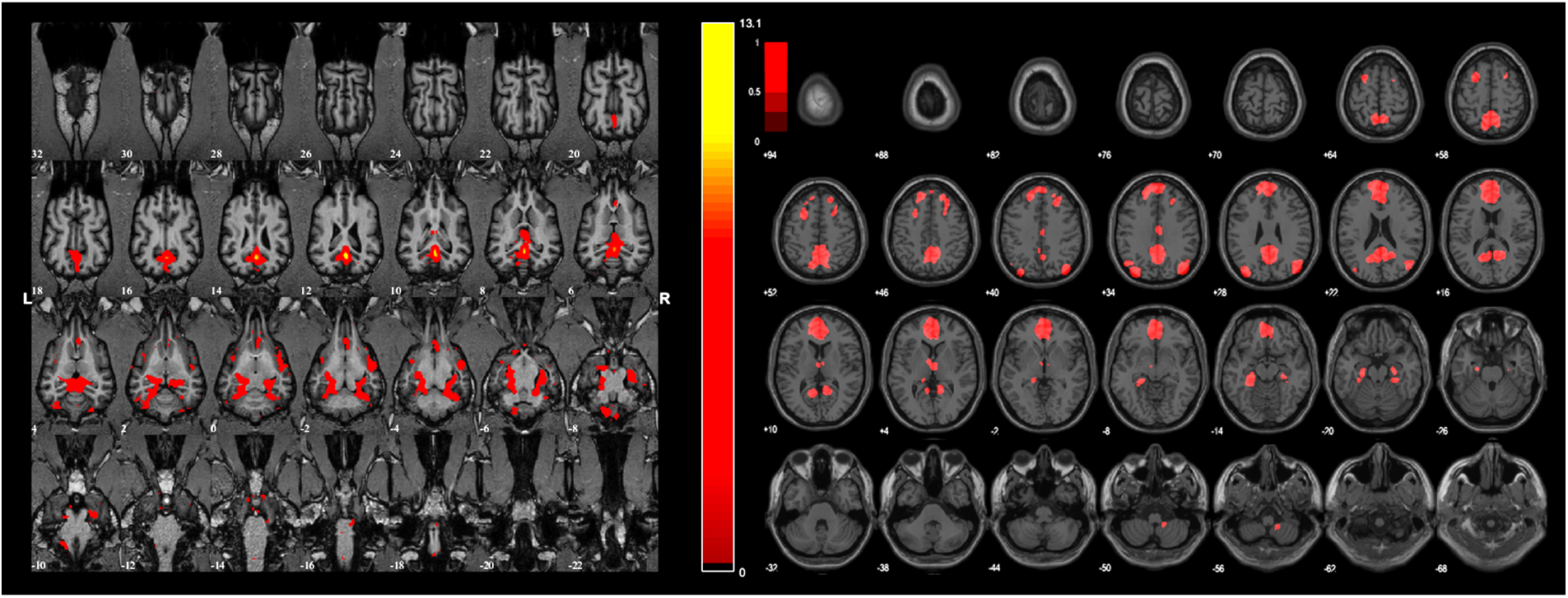
Left: RSN A, showing the dog default mode network (DMN). Overlaid color maps represent thresholded z-scores at a threshold of z>2. Right: Illustration of the human DMN from^37^.

**Table 1.**
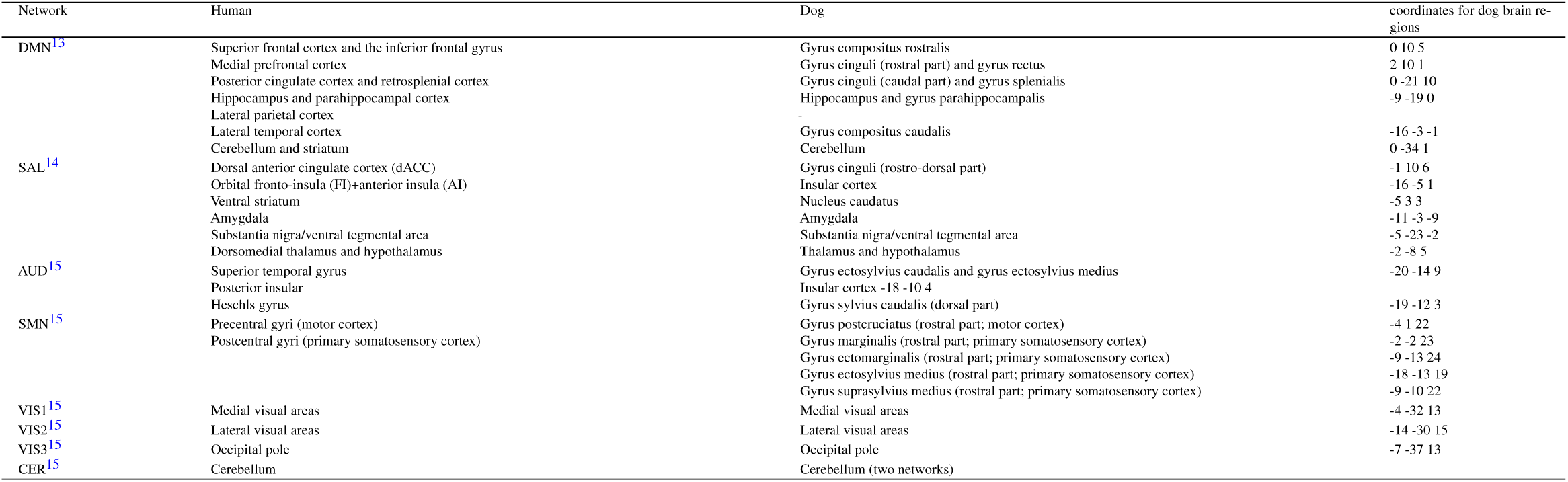
Brain regions involved in the respective human resting state networks based on the referred studies and the corresponding regions in the dog resting state networks. The dog brain regions coordinates refer to the stereotaxic, breed-averaged canine brain atlas^36^ In case of the default mode network, we found no involvement of lateral parietal regions in dogs.

RSN B (Fig. 2): This network contained the rostro-dorsal part of the gyrus cinguli, bilateral insular cortex, nucleus caudatus, amygdala, substantia nigra/ventral tegmental area, thalamus, hypothalamus and auditory cortices; namely the bilateral gyrus ectosylvius caudalis, gyrus ectosylvius medius, and dorsal parts of the gyrus sylvius caudalis. This component contains the regions involved in the saliency network (SAL) and the auditory network (AUD) as described in previous human studies^14, 15^. RSN C (Fig. 3): This network contained bilateral regions of the motor cortex (rostral region of the gyrus postcruciatus) and primary somatosensory cortex (rostral part of the gyrus marginalis; rostral part of the gyrus ectomarginalis; rostral part of the gyrus ectosylvius medius and rostral part of the gyrus suprasylvius medius). This component corresponds to the sensorimotor network (SMN) described previously in humans^15^.

**Figure 2.**
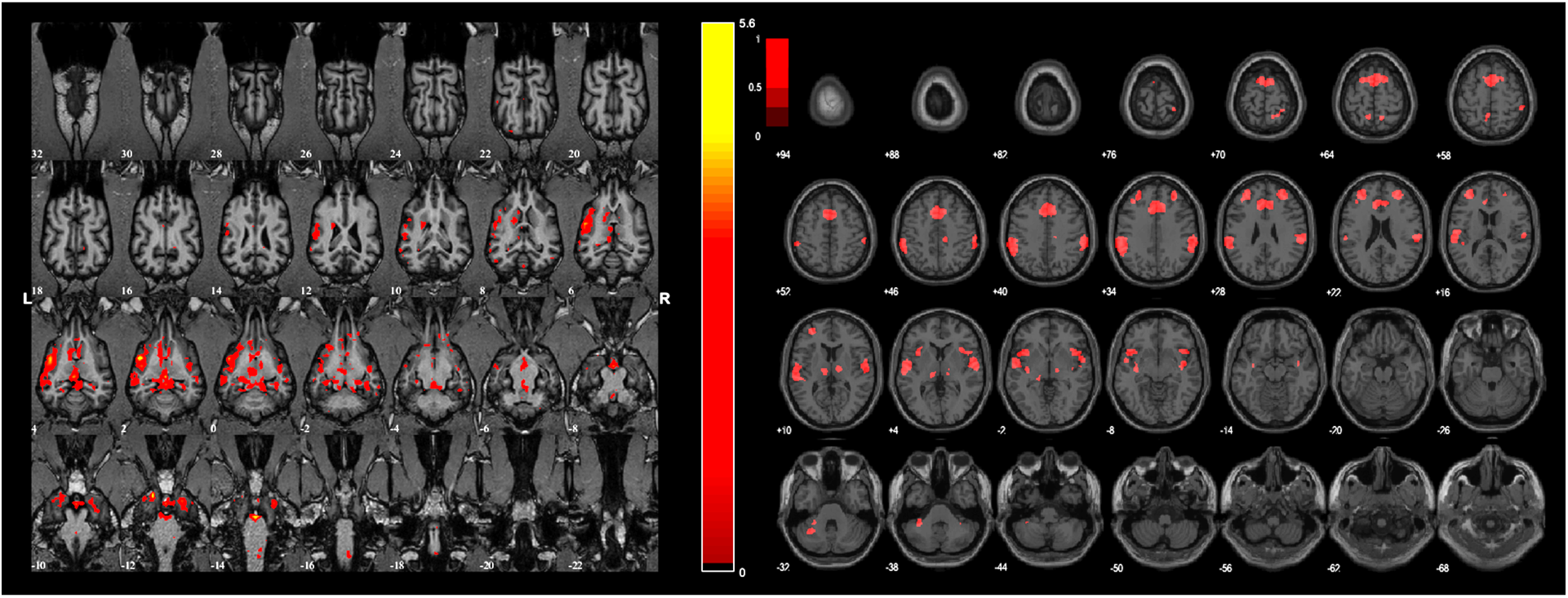
Left: RSN B, showing a combined saliency (SAL) and auditory (AUD) network in dogs. Overlaid color maps represent thresholded z-scores at a threshold of z>2. Right: Illustration of the human SAL and AUD network from^37^.

**Figure 3.**
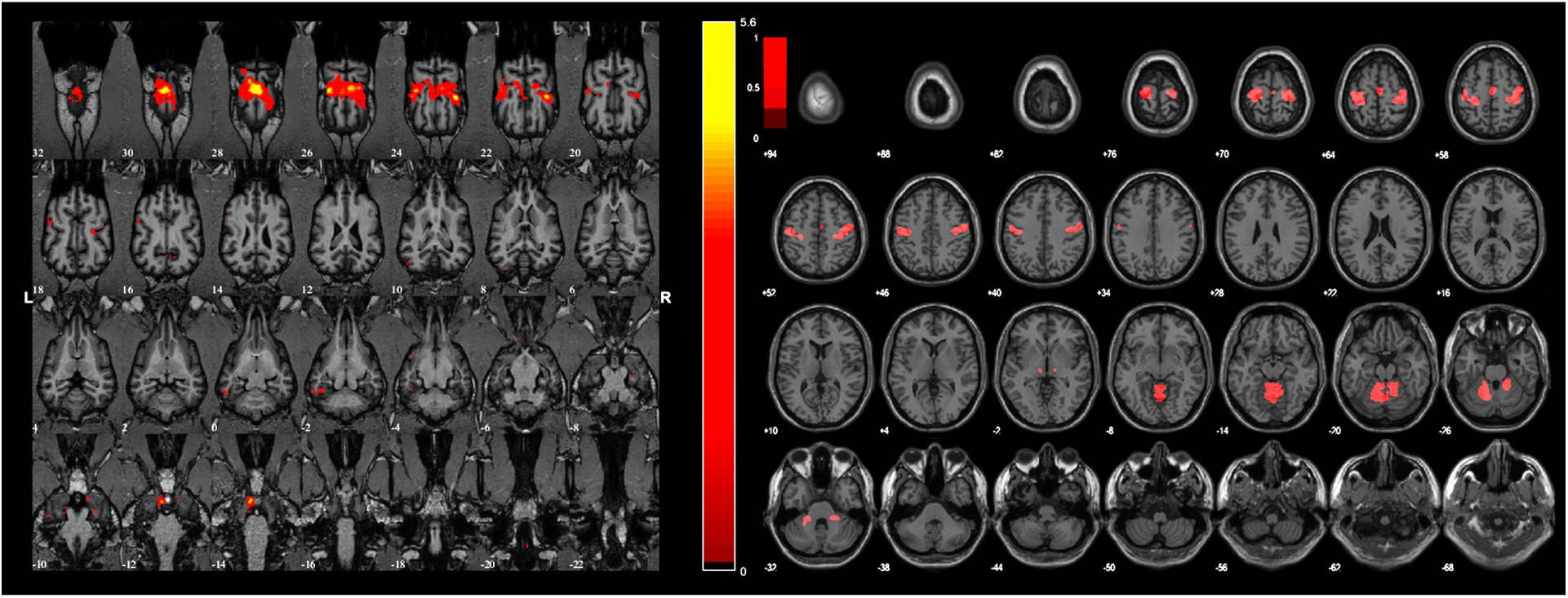
Left: RSN C, showing a sensorimotor network (SMN) in dogs. Overlaid color maps represent thresholded z-scores at a threshold of z>2. Right: Illustration of the human SMN network from^37^.

RSN D (Fig. 4): This network included the visual cortices, covering bilaterally the gyrus occipitalis, the gyrus suprasylvius medius, the gyrus ectomarginalis, the gyrus splenialis, caudal parts of the gyrus marginalis; and bilateral primary sensory areas, namely the gyrus postcruciatus, caudal regions of the gyrus marginalis, and middle region of the gyrus cinguli. This visual network contains regions corresponding to the human medial (VIS1) and lateral visual areas (VIS2) and the occipital pole (VIS3).

**Figure 4.**
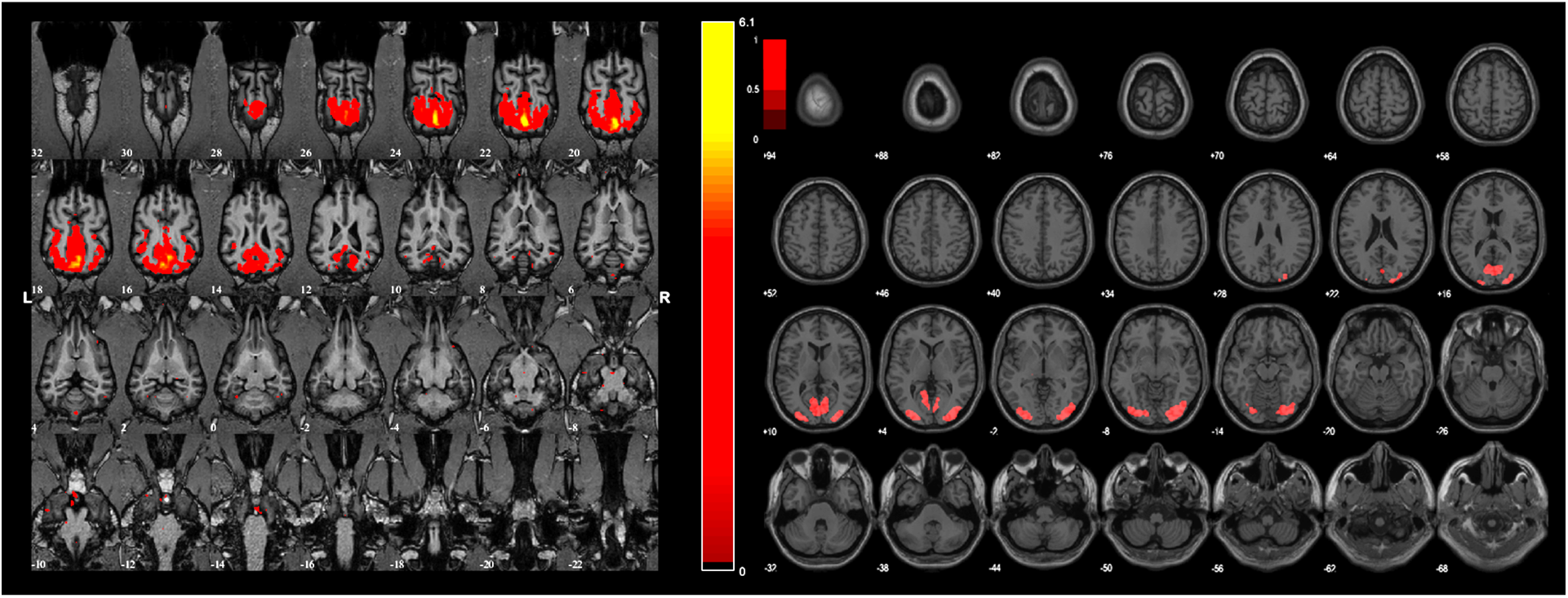
Left: RSN D, showing a visual network (VIS) in dogs. Overlaid color maps represent thresholded z-scores at a threshold of z>2. Right: Illustration of the human primary and higher visual network from^37^.

RSN E and RSN F (Fig. 5 & 6 Cerebellar network components, covering both hemispheres of the cerebellum and the cerebellar vermis, with RSN E located caudally, while RSN F involving more rostral parts of the cerebellum. This component is analogue to cerebellar networks (CER) previously reported in human^15^ and non-human animal studies^5, 7^.

Comparing the three investigated model dimensionalities revealed that the combined auditory-salience and cerebral network emerged at d=20 and at d=30, while the other networks (DMN, VIS, SMN) were present at all three investigated model orders.

**Figure 5.**
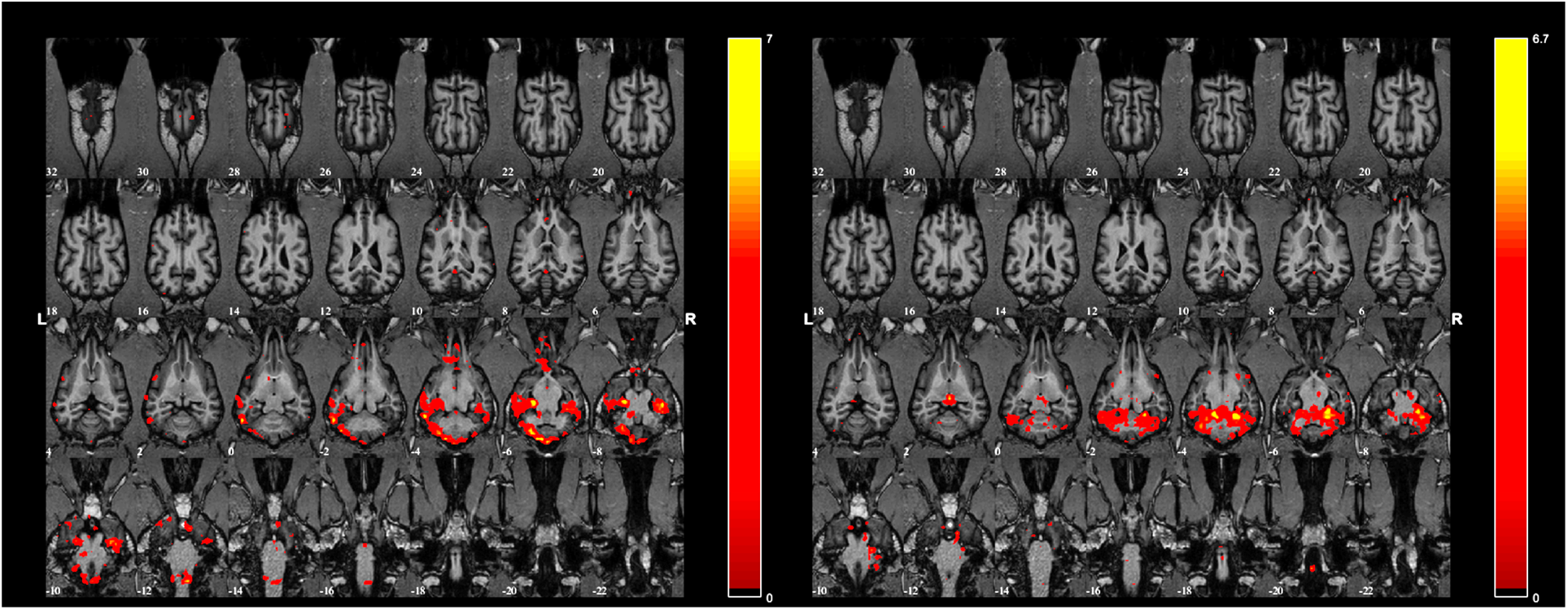
Left: RSN E. Right: RSN F. Two cerebellar networks in the dog brain. Overlaid color maps represent thresholded z-scores at a threshold of z>2.

## Discussion

We described multiple, spatially distributed RSNs in awake, unrestrained dogs, the evolutionarily most distant taxa from humans scanned without anesthesia so far. Our results indicate that dogs’ default mode network shows antero-posterior connectedness, containing areas from both the prefrontal cortex and the anterior cingulate cortex, with additional involvement of hippocampal regions, but it does not include the parietal cortex. In humans, these parietal regions (angular gyrus, temporoparietal junction) are thought to be involved in complex cognitive processes such as accessing conceptual representations about events or items and theory of mind^13^, cognitive functions whose extent and level in dogs are debated^16^. Our results do not support previous results^9^, however our study diverged in several aspects which could account for the different findings. We utilized a larger sample size (4 vs. 22), longer data acquistion runs, different temporal filtering and lower dimensionality in the ICA.

In our study, dogs salience network also contained the auditory cortex, unlike in anesthetized ferrets where a separate auditory network was found^10^, but similarly to previous findings in awake marmosets^7^. To understand if this fusion in dogs reflects real functional differences in auditory processing compared to humans, or if it is a consequence of dimensionality or differences e.g. in brain size, further studies are needed. Both in dogs and marmosets, the insular cortex is located very close to the primary auditory cortex and both regions are relatively small, which may hinder their split into separate, spatially independent components in this type of analysis.

We found a single visual network component, a finding similar to that of ferrets^10^, also a member of the order Carnivora. The presence of a single visual network falls between the lack of a separate visual network in rodents^17^ and the multiple visual networks found in primates^7^, which is in line with the ecology of the species. While the visual cortex of mice and rats is relatively small^18^ and this species rely little on vision, primates have a well-developed visual cortex and this taxon heavily relies on vision^19^. Dogs’ vision is inferior to that of primates, but they still rely on vision under a wide range of light conditions, they are visual generalists^20^. Dogs single visual network may dissociate into subnetworks in case of a higher number of extracted components, but this was not the case for the applied network dimensionalities in our study (d=20 and 30).

Dogs have some advantage regarding the scanning setup when compared to other model animals in fMRI studies. After special training, dogs are suitable to be scanned without sedation, which is beneficial as anaesthesia has a large impact on resting state functional connectivity^21^, and unlike the other non-human animal species scanned awake, there is no need of mechanical restraints either, which also have the risk of influencing rs-network connectivity^22^. The dog subjects are willingly motionless during scanning for an extended period of time (as performing a conscious action inhibiting their movements), a situation more closely resembling the conditions of human rs-fMRI measurements. While one may argue that the dogs are executing a rewarded “task” and not really resting, it is important to keep in mind that humans also consiously perform the same “hold still” task during fMRI measurments and most often do so in exchange to some predetermined reward (in form of financial compensation). Additionaly, while resting state networks are usually analysed in data collected during rest, they are also present when performing cognitive tasks^23^.

## Methods

### Subjects

We measured 22 family dogs *(Canis familiaris)* (age 6.41 *±* 3.42 years (mean *±* SD), range 2-13 years, 10 females and 12 males, 8 golden retrievers, 6 border collies, 1 Labrador retriever, 1 labradoodle, 2 mongrels, 1 Chinese crested dog, 1 Cairn terrier, 1 Hungarian vizsla, 1 Australian shepherd). Training procedure has been described in detail in a previous study^24^, and was based on individual and social learning using positive reinforcement.

### Experimental procedure

The experiment consisted of a 2-minute-long pretraining, to familiarize the dogs to the semi-continuous scanning procedure, and two 6-minute-long data collection runs. To provide sound protection, the dogs were wearing ear muffs. During scanning, dogs were lying with their eyes open, without presentation of a fixation cross, with their handler being visible, but avoiding eye contact with the subject. Motion threshold for successful runs was set to a maximum of 3 mm (for each translation direction) and 3 degree (for each rotation direction) during the whole run.

### Image acquisition

Functional MRI acquisitions were performed on a Siemens Prisma 3T scanner Siemens Healthcare, Erlangen, Germany) using a Gradient Echo Echo Planar Imaging (GRE-EPI) sequence with TR = 2640 ms including a 500-ms delay at the end of each volume, and TE = 30 ms. The protocol had an in-plane Field-of-View 128 mm x 128 mm, using 2 mm in-plane resolution and 2 mm slice thickness, measuring 31 slices with an inter-slice gap of 0.5 mm, using an excitation flip angle of 86?r. Phase-encoding direction was set to left-right. A single loop coil was used for signal detection, fixed onto the head of the dog and to the table of the scanner. In each run, 139 volumes were acquired, with the first 2 of them being discarded before processing, resulting in a total functional scanning time of 367 seconds/run. A T1-weighted anatomical scan was carried out separately as part of another study on each awake dog for spatial registration on a 3T Philips Ingenia scanner (Philips Medical Systems, Best, The Netherlands), using a 3D Turbo Field Echo (TFE) sequence, with TR = 9.85 ms, TE = 4.6 ms, and an isotropic resolution of 1 mm.

### Image analysis

FMRI preprocessing included affine realignment (6 parameters, least square aproach) and reslicing of the 137 images of the individual runs in SPM12 (http://www.fil.ion.ucl.ac.uk/spm/), followed by manual coregistration of the mean image to the individuals’ own structural T1 image in Amira 6.0 (Thermo Fisher Scientific). The structural image was normalized and transformed (linear, non-rigid transformation) to the template brain in Amira. The study-specific template was selected from the 22 individual structural images by a veterinary neurologist (K.C.), to ensure that the specimen is representative of our mesocephalic sample in regard to shape, size, gyrification and is free from any detectable deformity. The resliced images were then coregistered and normalized to this transformed mean functional image via SPMs standard nonlinear warping function with 16 iterations.

We applied high pass filtering^25^ with a 0.01 Hz cutoff in MATLAB 2017 environment (Mathworks, Sherborn, MA). We decided against censoring via removing volumes midrun from our functional datasets, because censoring in combination with frequency filtering can intoduce additional artefacts^26^ and excessive motion was not prevalent (scan to scan displacement larger than 0.2 mm occured in 5 % of the scans). Two runs had been censored due to exceeding motion threshold, after 130 and 133 scans respectively (out of 137). Runs with supra-threshold motion occurring before the 130th scan (exceeding 3 mm in any direction or more than 3?r rotation in any direction during the total length of the run) were repeated. The average scan-to-scan displacement was below 0.04 mm for each translation direction, and below 0.0007 degree for each rotation direction in case of all runs. The mean maximal scan to scan movements per run (averaged across all runs and subjects) was below 0.77 mm for each translation direction, and below 0.01 degree for each rotation direction. Therefore, censoring of fMRI volumes during runs due to excessive motion, as applied by e.g.^27^ was deemed inappropriate, because only 3 volumes (each in separate runs) out of the 5975 collected images during the study (0.05 %) would have been affected by this treshold.

We applied a whole dog brain mask without segmentation of white matter or cerebrospinal fluid, as these parts are informative during evaluation of the components. If a component involves a large number of small clusters mainly located in the white matter, cerebrospinal fluid or near to blood vessels (particularly arteries) it usually suggests that the component’s origin is physiological noise (respiration, pulsation)^28^. After removal of the mean per time point a standard, Spatial Group Independent Component Analysis was carried out via GroupICATv4.0b (GIFTv3.0b, Medical Image Analysis (MIA) Laboratory) with the following settings: subject-specific spatial PCA with Infomax GICA back reconstruction, with a serial group analysis, number of components set to 20. As the choice of model order can have a large effect on ICA decompositions, to evaluate the robustness of the ICA-identified maps, we performed additional exploratory analyses with models specified by 10 and 30 components as well. This showed that the dimensionality of 20 component still yielded stable, bilateral signal components in recognisable neuroanatomical systems^29^ while the described signal components were also present in case of 10 and 30 components. Subject Specific PCA is done on each data-set before doing group ICA and GICA was shown to be a more robust tool to back reconstruct components for low model order^30^. To evaluate component stability, ICASSO stability analysis was performed (number of iterations = 10) and only components with Iq>0.8 were considered^31^.

Components were converted to Z scores, and a threshold of Z>2 was applied. We chose to use Z>2 in order to slightly exceed treshold of similar exploratory studies in other species (e.g. mice^5^, rat^32^ and ferret^10^). Classification of signal components was based on spectral characteristics (power spectra, dynamic range, LF to HF power ratio) and evaluation of the spatial maps following the guidelines provided by^33^ and^28^. Signal components are indicated by higher dynamic range and LF to HF power ratio, an oscillatory time course, with a low number of large clusters located mainly in the grey matter, following known anatomical boundaries. Components showing characteristics of susceptibility, motion, vascular of ventricular artefacts were excluded from further analysis. Anatomical labelling was carried out based on relevant anatomical brain atlases^34–36^.

## Acknowledgements

This project has received funding from the European Research Council (ERC) under the European Unions Horizon 2020 research and innovation programme (Grant Agreement No. 680040), was supported by the National Research, Development and Innovation Office (Grant No. 115862K) for MG, the Hungarian Academy of Sciences [MTA-ELTE Comparative Ethology Research Group (Grant No. F01/031), MTA-ELTE Lendület Neuroethology of Communication Research Group (Grant No. 95025), the János Bolyai Research Scholarship of the Hungarian Academy of Sciences for EK and AA], and the Eötvös Loránd University. ÁK was supported by the Hungarian Brain Research Program (Grant No. 2017-1.2.1-NKP-2017-00002) and ÁM received support from the Program of National Excellence (NKPÓ17, 2017-1.2.1-NKP-2017-00002). We thank all dog owners participating in our study.

## Author contributions statement

D.SZ. conceived the experiment; D.SZ., Á.K., M.G. conducted the experiment; D.SZ., K.C. and Á.K. analysed the results. All authors reviewed the manuscript.

## Additional information

Link to multimedia model illustrating the dog resting state networks as a 3D composite image: (https://www.youtube.com/watch?v=YMxNNIfPkBs)

